# *In situ* structural basis for the LHCSR3-mediated photoprotection of photosystem II

**DOI:** 10.64898/2026.07.02.736019

**Authors:** Xiaodong Su, Chunling Wu, Shengrong Cui, Zhenfeng Liu, Xinzheng Zhang, Mei Li

## Abstract

Energy-dependent quenching (qE) represents a critical mechanism for photosynthetic organisms to mitigate photodamage caused by excessive light. In green algae, LHCSR3 protein plays a central role in qE, activated by thylakoid lumen acidification and associated with photosystem II (PSII) to dissipate excess energy. Despite extensive efforts, the assembly and energy dissipation mechanisms of PSII-LHCSR3 have remained unknown. Here, we present the *in situ* structures of the PSII supercomplex embedded in the native thylakoid membranes of *Chlamydomonas reinhardtii* in both quenched (LHCSR3-bound) and unquenched (LHCSR3-free) states at near-atomic resolutions. Our results demonstrate that in high-light-acclimated cells, LHCSR3 binds to PSII peripheral antenna CP26 and associates with an extra LHCII trimer (eLHCII). Structural comparison of LHCSR3 with other light-harvesting complexes reveals possible protonation-induced conformational changes in LHCSR3 and rearrangements of its two pigment clusters, which potentially serve as quenching sites to dissipate excess energy transferred from CP26 and eLHCII. Our findings provide a direct visualization of how photoprotection is spatially organized *in vivo* and have implications for engineering natural and artificial photosynthetic systems with more dynamic photoprotection.

## INTRODUCTION

Solar energy is indispensable for photosynthesis, yet excess light energy can cause oxidative damage to photosynthetic complexes ^1^. Photosystem II (PSII) is the most vulnerable one among various photosynthetic complexes and the primary target of photodamage ^2^. To avoid or reduce photodamage, plants and algae have developed sophisticated photoprotective mechanisms to safely dissipate excess energy, a process known as nonphotochemical quenching (NPQ) ^3^. Among various components of NPQ, the energy-dependent quenching (qE), induced and relaxed on a timescale of seconds to minutes, represents the most rapid and major photoprotective strategy adopted by plants and algae ^2,4,5^. The qE process is triggered by thylakoid lumen acidification, which results from the increased accumulation of protons caused by the accelerated photosynthetic electron flow under excess light conditions ^6,7^. In plants and green algae, qE is mainly mediated by the PSII subunit S (PsbS) ^8^ and the Light Harvesting Complex Stress Related (LHCSR) protein ^9^, respectively. Both proteins have been suggested to be activated via sensing lumen acidification and associate with PSII–light-harvesting complex II (PSII-LHCII) supercomplexes to induce qE ^10,11^.

LHCSR is a member of the LHC superfamily with three transmembrane helices (TMHs) and binds chlorophyll and carotenoid molecules, potentially acting as a direct quencher to dissipate excess energy using its pigments ^12^. The expression of LHCSR is upregulated under high-light conditions, and depletion of LHCSR significantly reduces NPQ in *Chlamydomonas reinhardtii* (*C. reinhardtii*, a model organism of green algae) ^13^. *C. reinhardtii* possesses three LHCSR isoforms, namely LHCSR1, LHCSR3.1 and LHCSR3.2. The two LHCSR3 isoforms have identical amino acid sequences of their mature proteins, while LHCSR1 and LHCSR3 share 82% sequence identity ^9^. Previous studies have demonstrated that LHCSR3 plays a major role in NPQ of *C. reinhardtii*, while LHCSR1 partially compensates for the loss of LHCSR3 in the *npq4* mutant (*lhcsr3* gene knockout) and also contributes to NPQ ^9,11,13^. Three acidic amino acid residues located in the lumenal regions of LHCSR3 play a key role in sensing the decrease of lumen pH through protonation, and the triple mutant exhibits a 72% reduction in qE levels *in vivo* ^14,15^. *In vitro* refolding of recombinant LHCSR3 protein with pigment molecules suggested that LHCSR3 binds 6-8 chlorophyll molecules, mostly chlorophyll *a* (Chl *a*), and 2-3 carotenoids, presumably one lutein (Lut) and one violaxanthin (Vio) ^12,16^. Vio belongs to the xanthophyll cycle carotenoids and its conversion to zeaxanthin (Zea) under high-light plays a crucial role in NPQ ^17,18^. While it was suggested that Zea is dispensable for NPQ in *C. reinhardtii* ^11,19^, a recent work demonstrated that Zea-bound LHCSR3 exhibits a shorter fluorescence lifetime than the Vio-bound form and an obvious quenching can be observed in the *zep* strain (Zea-enriched) even at neutral pH ^20^. Therefore, two distinct quenching mechanisms of LHCSR3, depending on either pH or Zea, are likely operating in *C. reinhardtii*.

While earlier reports have suggested the presence of multiple potential LHCSR3-binding sites in the peripheral regions of PSII, including one nearby LhcbM1 of the LHCII trimer ^21,22^ and the other adjacent to the PSII core subunit PsbR ^23^. Moreover, two minor LHCs, namely CP26 and CP29, were previously shown to be involved in energy quenching and may also serve as interaction partners of LHCSR ^24,25^. Analysis of the two-dimensional negative-stain projection maps of PSII suggested that LHCSR3, in either dimeric or monomeric form, binds to C_2_S_2_ type (C: PSII core; S: strongly-bound LHCII, S-LHCII) PSII-LHCII at multiple positions close to the S-LHCII trimer and CP26 ^26^. However, the results are inconclusive as the projection maps at low resolution are insufficient for unambiguous assignment of LHCSR3. Despite extensive investigations of LHCSR3 and the availability of numerous biochemical and spectroscopic data, the mechanistic understanding of LHCSR3 in qE remains limited, partly due to the lack of structures of LHCSR3 and PSII-LHCSR3 complex. Fundamental questions regarding how LHCSR3 is activated and how it binds to PSII to quench excess energy remain to be addressed. Here, we present the *in situ* structures of the PSII-LHCII with and without bound LHCSR3, from *C. reinhardtii* cells acclimated under high-light and low-light conditions respectively. The structures represent the first high-resolution *in situ* observation of PSII-LHCII supercomplexes in both quenched and unquenched states, providing detailed information on the structure of LHCSR3 and its binding mode with PSII, and revealing potential quenching mechanisms underlying the LHCSR3-mediated photoprotection of PSII.

## RESULTS

### Capturing LHCSR3-bound PSII *in vivo* using newly developed *in situ* methods

*C. reinhardtii* CC124 cells were cultured under continuous low-light of 30-40 μmol photons m^-2^ s^-1^ (LL-cells). To induce quenching, LL-cells were further treated with high-light of 500 μmol photons m^-2^ s^-1^ (HL-cells). Consistent with previous results ^9,12^, an obvious increase of LHCSR3 expression and Zea accumulation in HL-cells were observed when compared with LL-cells (Fig. S1A-C). Furthermore, our NPQ analysis showed that the NPQ amplitude in HL-cells is substantially higher than that in LL-cells, and the PSII activity remains largely unaffected (Fig. S1D, E), suggesting that PSII is effectively protected by LHCSR3 and/or Zea molecules in HL-cells and is presumably maintained in a quenched state.

Both HL- and LL-cells were deposited onto grids and subsequently vitrified. The HL conditions were maintained throughout the HL-grids preparation to preserve the quenched state. Cryo-lamellae were prepared on both HL- and LL-grids using cryo-focused ion beam (cryo-FIB) milling (Fig. 1A). Furthermore, to minimize damage to the lamellae caused by cryo-FIB milling, we developed cryogenic low-energy polishing in FIB (cryo-LEP-FIB) technique ^27^, which greatly improves the quality of thinned lamellae. The cryo-electron microscopy (cryo-EM) data were subsequently collected from these lamellae. *In situ* particle picking of PSII complexes is highly challenging as they are mostly embedded in thylakoid membranes with strong background. To overcome this challenge, we utilized the *in situ* single-particle analysis (isSPA) method ^28,29^ to locate various PSII complexes in the native thylakoid membranes using the structure of dimeric PSII core (C_2_) as the searching template (Figs. S2, S3). Finally, we obtained the reconstructions of the C_2_S_2_M_2_L_2_–type (M/L: moderately-/loosely-bound LHCII, M-LHCII/L-LHCII) PSII-LHCII supercomplexes in HL- and LL-cells at resolutions of 3.1 Å and 3.4 Å, respectively (Fig. 1, Figs. S2, S3, Table S1). We found that the C_2_S_2_M_2_L_2_ complex represents the major type of PSII-LHCII in *C. reinhardtii*, consistent with previously reported cryo-electron tomography analysis of *C. reinhardtii* thylakoid membranes ^30^.

**Figure 1.**
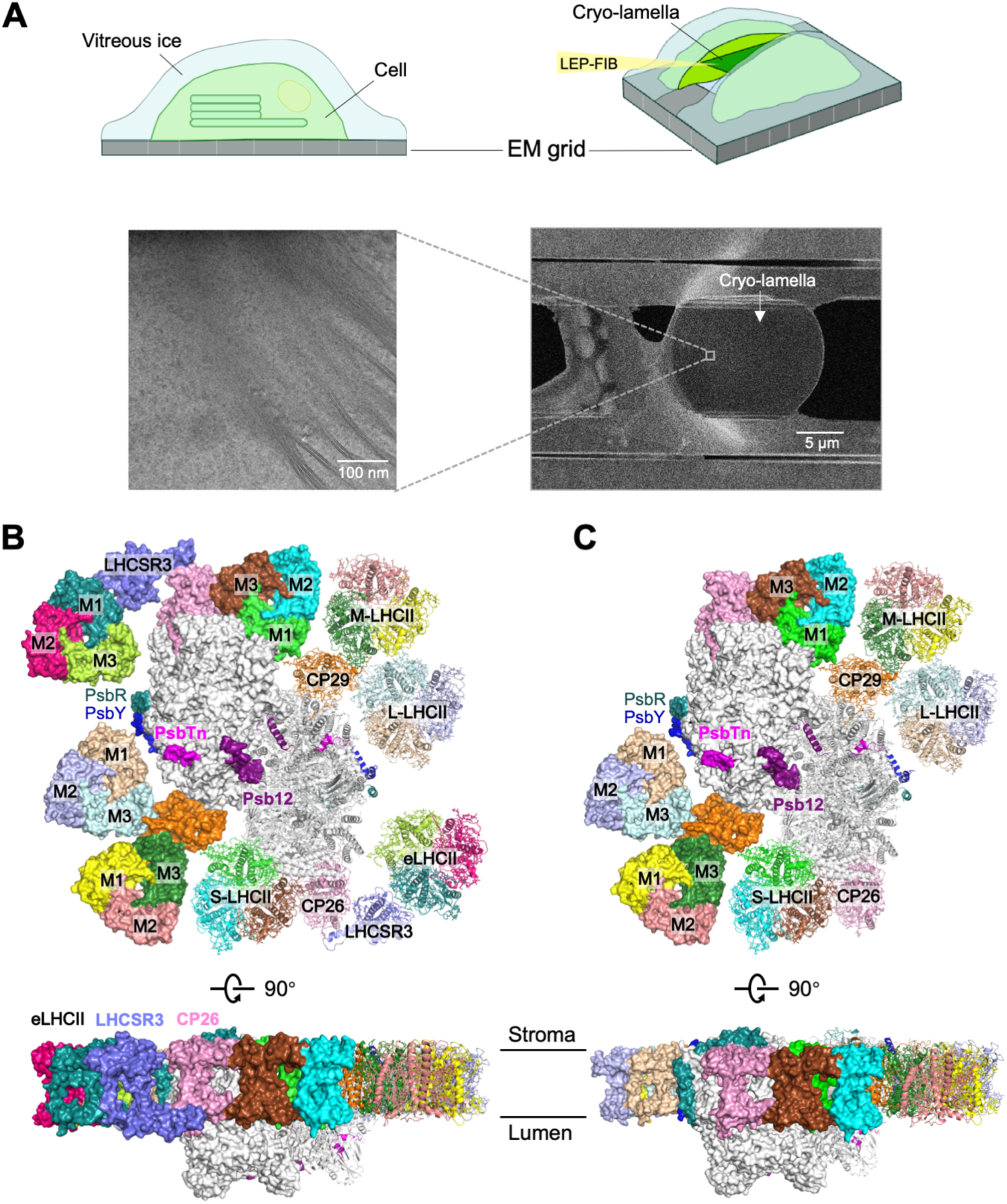
*In situ* structure determination of HL_PSII-LHCSR and LL_PSII. **(A)** A schematic diagram of cryo-cellular lamella preparation and a representative image of the cryo-lamella. **(B-C)** Overall structure of HL_PSII-LHCSR (B) and LL_PSII (C) shown in cartoon and surface mode for each monomeric moiety. Light-harvesting antennae and four core subunits (Psb12, PsbR, PsbY and PsbTn) are shown in different colors and labeled. Other PSII core subunits are shown in white.

Strikingly, one monomeric and one trimeric LHC proteins were observed at the peripheral regions of both PSII monomers in the C_2_S_2_M_2_L_2_ supercomplex acclimated to HL-conditions, and they are absent in the LL-adapted complex (Fig. S4). Focus refined density map enables us to identify them as one LHCSR3 monomer and one LHCII trimer (Fig. S5), binding to the site nearby CP26 (Fig. 1B). We thus named the *in situ* structures of PSII supercomplexes from HL- and LL-cells as HL_PSII-LHCSR (HL-structure) and LL_PSII (LL-structure), respectively. It is evident that the dimeric PSII-LHCII supercomplex exists by itself at low-light, whereas under high-light conditions, the PSII complex becomes associated with a LHCSR3 monomer and an extra LHCII trimer (eLHCII). These structural observations suggest that the HL_PSII-LHCSR and LL_PSII structures represent the complexes in a quenched state and a light-harvesting state, respectively. Under high-light condition, binding of LHCSR3 to CP26 establishes a close connection between LHCSR3 and PSII-LHCII, and LHCSR3 further associates with the eLHCII at a newly discovered peripheral site of the complex.

Except for the presence/absence of the LHCSR3-eLHCII subcomplex nearby CP26, the C_2_S_2_M_2_L_2_ moiety in HL- and LL-complexes exhibit no evident conformational differences (Fig. S6). While minor conformational changes are observed in amino acid side-chains and pigment molecules, they could not be unambiguously distinguished from structural errors at the current resolutions. Nevertheless, our reconstructions allow us to identify all three major LHCIIs in the C_2_S_2_M_2_L_2_ moiety as LhcbM1-LhcbM2-LhcbM3 heterotrimers (Fig. S7), and build relatively complete structural models of several core subunits, including PsbTn, PsbR and PsbY (Fig. 1B, C, Fig. S8, Table S2). In addition, we identified an uncharacterized protein (XP_001690252.1) located at the lumenal interface between adjacent PSII monomers in both structures, and named it Psb12 (Fig. 1B, C, Fig. S9). Psb12 interacts with CP47 from one PSII monomer and D1’/PsbO’ from the adjacent monomer, thus contributing to the stability of the PSII dimer (Fig. S9E). Apparently, PSII complexes are structurally more intact and contain more loosely-associated peripheral proteins under the *in vivo* conditions than the *in vitro* purified ones ^31,32^, underscoring the necessity and advantages of using *in situ* approaches to capture the LHCSR3-bound PSII supercomplexes in their native conformational states.

### The *in situ* structure of LHCSR3

The LHCSR3 protein adopts a typical LHC fold, possessing three TMHs, termed helices B, C and A from the N- to the C-terminus, as well as loops connecting TMHs B and C (BC loop) and TMHs C and A (AC loop) (Fig. 2A, B). In addition, LHCSR3 features an elongated C-terminal domain at the lumenal side, consisting of a long α-helix (helix D) followed by an irregular loop. This structural feature is unique to LHCSR, and has not been observed in other LHC proteins. In LHCSR3, we identified eight chlorophylls and two stably-associated carotenoid molecules at the central region (Fig. 2C-F), consistent with previous analysis of the pigment-binding property of the recombinant LHCSR3 protein ^12,16^. Remarkably, we also observed two extra polyene-like densities at the peripheral region of LHCSR3 (facing eLHCII), which fit well with two carotenoid molecules (Fig. 2F).

**Figure 2.**
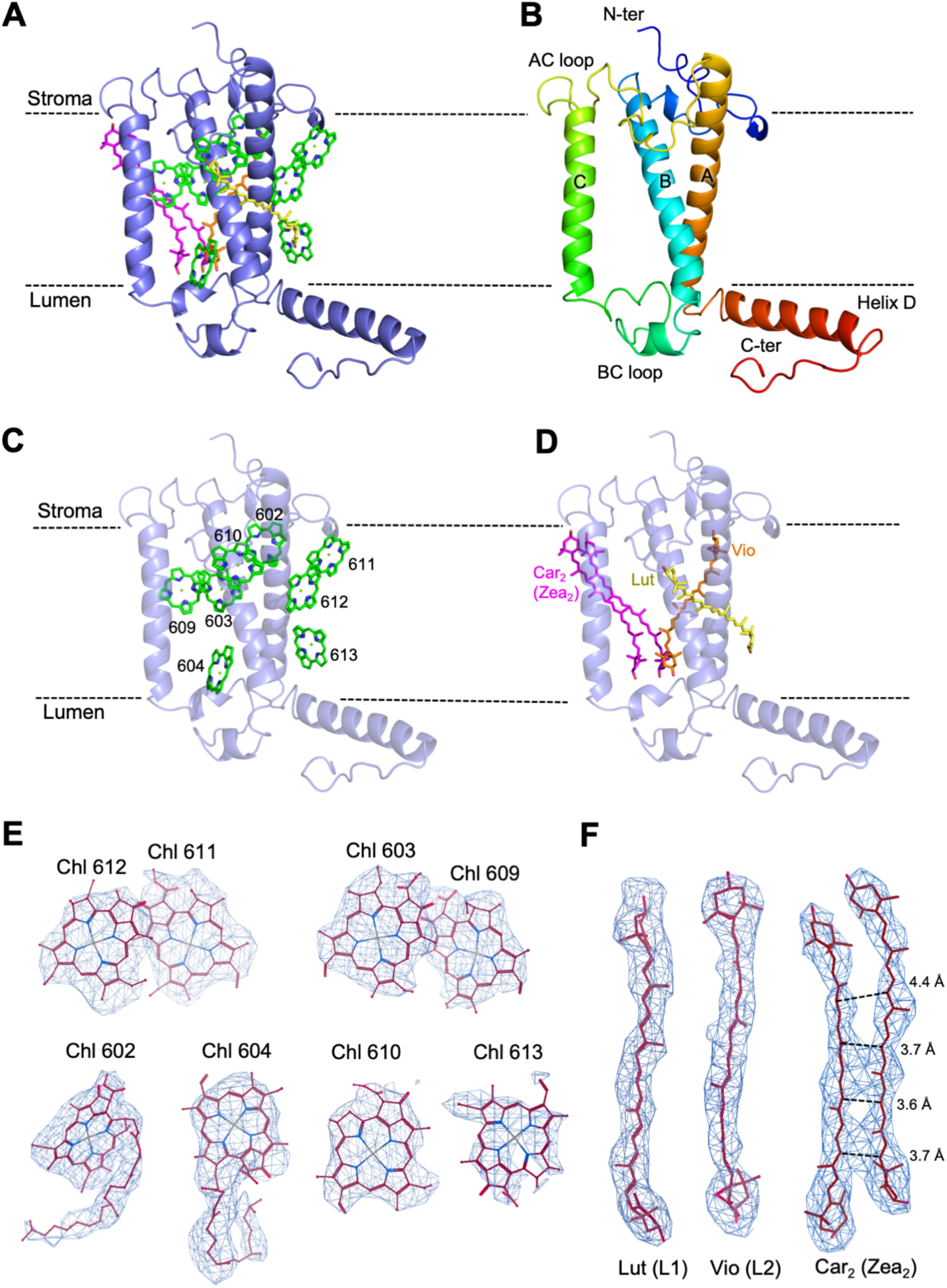
*In situ* structure of LHCSR3. **(A)** Cartoon representation of LHCSR3. Chlorophyll *a*, lutein, violaxanthin and the carotenoid dimer (potentially a Zea dimer) are shown as green, yellow, orange and magenta sticks respectively. The phytol tails of chlorophylls are omitted for clarity. **(B)** LHCSR3 apoprotein structure colored in rainbow mode. Three transmembrane helices are labeled as B, C and A. The loops connecting helix B and C and connecting helix C and A are labeled as BC loop and AC loop, respectively. The helix D, N-terminus (N-ter) and C-terminus (C-ter) are labeled. **(C, D)** Arrangement of chlorophyll (C) and carotenoid (D) molecules in LHCSR3. The two carotenoid molecules are tentatively assigned as a Zea dimer and is labeled as Car_2_ (Zea_2_). **(E, F)** Cryo-EM densities of eight chlorophylls (E) and four carotenoids (F) in LHCSR3.

All eight Chl binding sites found in LHCSR3 are conserved in LHCII, with six chlorophylls (Chls 602, 603, 609, 610, 612, 613) coordinated by amino acids identical to those in LHCII (represented by LhcbM1) (Fig. S10). The remaining two chlorophylls (Chls 604 and 611) are typically coordinated by a water and a phosphatidylglycerol (PG) molecule, respectively, in most other LHCs. However, LHCSR3 lacks this Chl-coordinating lipid molecule, presumably due to the substitution of an asparagine (N199) in LHCSR3 for the PG-stabilizing lysine in other LHCs (Fig. S11). Therefore, both Chls 611 and 604 in LHCSR3 are likely coordinated by water molecules, which were not resolved in our 3.1 Å resolution map. It is noteworthy that Chl 613 exhibits weaker density compared to other chlorophyll molecules (Fig. 2E), indicating that under strong light, the chlorophyll only loosely binds to or partially occupies the 613 site in LHCSR3. We tentatively assigned all eight chlorophyll molecules as Chl *a*, as previous studies have demonstrated that LHCSR3 has strong affinity for Chl *a*, and the lack of Chl *b* does not affect its proper folding ^12^.

The two stably-associated carotenoids in LHCSR3 are located at the central L1 and L2 sites, which are conserved in LHCs and have crucial roles in the folding and structural stability of LHC proteins ^33,34^ (Fig. S10C). Based on previous biochemical and *in vitro* reconstitution results ^12,35^, we assigned one Lut at L1 site and one Vio at L2 site. In addition, we found two loosely-bound carotenoids at a novel peripheral site of LHCSR3 (Fig. 2D). These two carotenoid molecules are located in close proximity and almost parallel to each other, forming a tightly-coupled carotenoid dimer, which has not been observed in other LHC proteins. A previous study has shown that Zea and Vio exhibit specific tendency to form dimers (or aggregates) in detergent micelles ^36^. Computational modeling results indicate that the intermolecular distance of Zea dimers is approximately 3.7 Å ^37^, which agrees well with the distance between the two carotenoid monomers observed in our structure (Fig. 2F). In the HL_PSII-LHCSR complex, the carotenoid dimer is located at the outermost region exposed to the lipid bilayer (Fig. S12A), which is consistent with the proposed binding environment of xanthophyll-cycle carotenoids in protein complexes ^38^. Such an exposed location may enable Zea/Vio molecules to be readily accessed and converted by the zeaxanthin epoxidase/violaxanthin de-epoxidase ^39^ under low-light/high-light conditions. These analyses together suggest that the two carotenoid molecules may be a Zea dimer bound to LHCSR3 under high light. Our pigment analysis results further support this assignment, showing that while Zea and antheraxanthin (Ant) accumulate in HL-cells (Fig. S1C), the PSII-LHCII (mostly C_2_S_2_) and PSI-LHCI supercomplexes purified from HL-cells contain almost no Zea/Ant molecules (Fig. S12B, C), implying that Zea molecules do not assemble with these supercomplexes but are rather located outside of them ^38^. Based on these results, we tentatively assigned this carotenoid dimer as a Zea dimer (Fig. 2D), which is stabilized by several conserved residues in the AC and BC loops of LHCSR3 (Fig. S12D). Furthermore, we also observed a tube-shaped density near helix A at the stromal side, which matches well with one half of a carotenoid molecule (Fig. S12E, F). This putative carotenoid is stabilized at one end through forming a hydrogen bond with N199 of LHCSR3, while the other end may exhibit high mobility due to the lack of interactions with the complex. However, given that it exhibits only partial density in our reconstruction, we did not model this putative carotenoid molecule in our structure.

### Potential protonation-induced conformational changes in LHCSR3

It is generally believed that LHCSR3-dependent qE is achieved through conformational switch of LHCSR3 from an unquenched state to a quenched state, induced by the decrease in thylakoid lumen pH ^14^. While our *in situ* structure of LHCSR3 may represent its quenched conformation, the unquenched conformation remains unclear. Diatoms possess Lhcx proteins that are homologs of green algal LHCSR, and diatom Lhcx1 is suggested to be involved in quenching process ^40–42^. Previously, a structure of the PSII complex from diatom cells cultured under low light (light intensity of ∼50 μmol photons m^-2^ s^-1^) conditions was resolved ^43^. This complex contains a Lhcx6_1 monomer that is likely in its light-harvesting state. Therefore, we compared the sequences and structures of LHCSR3 with Lhcx6_1 and LhcbM1 (a typical Lhcb protein) to delineate potential conformational changes of LHCSR3 between its quenched and unquenched states (Fig. 3A-C, Fig. S10). We found that LHCSR3 differs considerably from other LHC proteins in the lumenal parts, with the most striking difference occurring in helix D, which pivots by approximately 60 degrees in LHCSR3 compared to that in Lhcx6_1/LhcbM1 (Fig. 3A, C). Moreover, BC loops in the three LHC proteins exhibit distinct conformation, with more pronounced difference between LHCSR3 and other LHCs located in the N-terminal half (BC-loop-N) (Fig. 3C). In contrast, the stromal parts of the three proteins are largely conserved, except that the N-terminal half of AC loop (AC-loop-N) in LHCSR3 and LhcbM1 contains an insertion which is absent in Lhcx6_1 (Figs. 3B, S10A). In LHCSR3, the extended loop formed by AC-loop-N contributes to the stabilization of the carotenoid dimer (Fig. S12D). Furthermore, the tilting angle of helix C differs in the three proteins (Fig. 3A).

**Figure 3.**
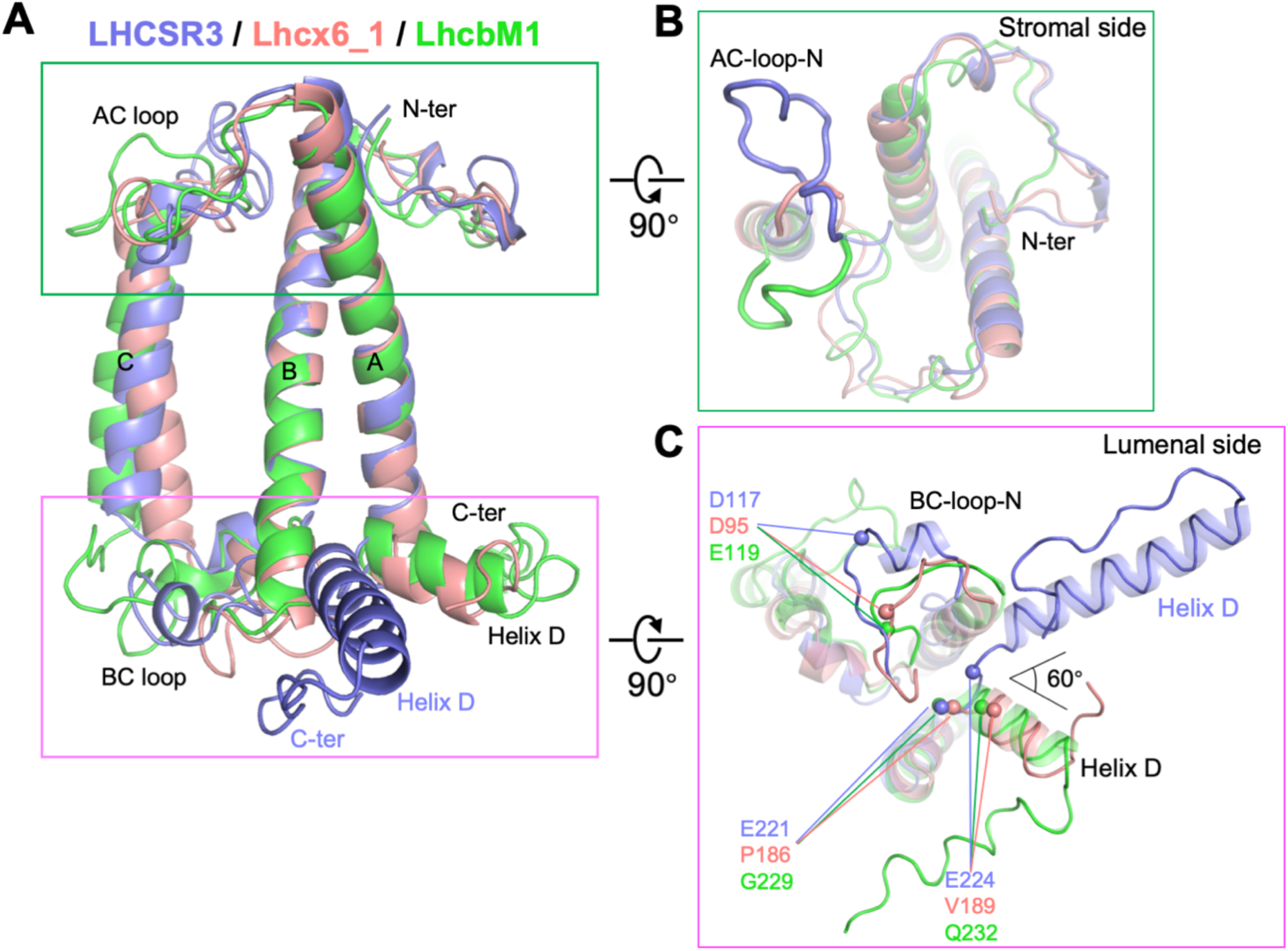
Potential protonation-induced conformational rearrangements of LHCSR3. **(A)** Structural comparison of *C. reinhardtii* LHCSR3 (slate) with diatom Lhcx6_1 (salmon, PDB code: 8IWH) and *C. reinhardtii* LhcbM1 (green). Proteins are shown as cartoon, and secondary structures are labeled. Pigment molecules are omitted for clarity. **(B)** Structural comparison of LHCSR3 with Lhcx6_1 and LhcbM1 viewed from the stromal side. Proteins are shown as transparent cartoon, with their N-terminal half of AC loop (AC-loop-N) also shown in ribbon mode and labeled. Both LHCSR3 and LhcbM1 are modeled with longer N-terminus than Lhcx6_1, and these N-terminal residues in LHCSR3 and LhcbM1 are omitted for clarity in (A) and (B). **(C)** Structural comparison of LHCSR3 with Lhcx6_1 and LhcbM1 viewed from the lumenal side. Proteins are shown as transparent cartoon, with their N-terminal half of BC loop (BC-loop-N) and C-terminal domain including helix D also shown in ribbon mode. Three acidic residues crucial for qE in LHCSR3 (D117, E221 and E224) and their corresponding residues in Lhcx6_1 and LhcbM1 are shown as spheres at their α-carbon atom positions and labeled.

Earlier studies revealed that three lumenal acidic residues, namely D117, E221 and E224, are crucial for pH-sensing and the LHCSR3-dependent NPQ induction ^15,44^. Notably, all three residues are located in regions where LHCSR3 differs in conformations from other LHC proteins (Fig. 3C, Fig. S10A). D117 is positioned in BC-loop-N region, which contains a short helix in LHCSR3, while the corresponding portion in other LHCs forms an irregular loop. E221 and E224 in LHCSR3 are situated in the loop region preceding helix D, whereas the residues equivalent to E224 in other LHCs are located inside helix D (Fig. 3C). This discrepancy is because helix D in LHCSR3 is one turn shorter at its N-terminus than that in other LHC proteins, resulting in a longer loop between the transmembrane domain and helix D in LHCSR3. This structural feature likely endows helix D with enhanced mobility, facilitating its large-angle rotation. Previous studies have indicated a correlation between conformational pH switch of lumenal helices in LHC proteins and the activation of quenching ^45,46^. Consistently, our structural comparisons suggest that protonation of the three qE-crucial residues in LHCSR3 may lead to conformational rearrangements of its BC-loop-N and the N-terminus of helix D, and a subsequent large-scale pivot of helix D.

### Specific interactions of LHCSR3 with CP26 and eLHCII

Earlier studies indicated that the NPQ activity is dramatically reduced in CP26-depleted mutant strains of *C. reinhardtii* and *P. patens* ^24,25,47^. In addition, previous fluorescence decay–associated spectral data revealed that LHCSR3 selectively inhibits the transfer of energy to CP43 ^48^, which primarily receives excitation energy from CP26 and S-LHCII. Our *in situ* HL_PSII-LHCSR structure explains these previous results, demonstrating that CP26 in the C_2_S_2_M_2_L_2_ supercomplex specifically mediates the binding of LHCSR3, and this binding depends on the unique orientation of helix D of LHCSR3 in quenched state. Under high-light conditions, the helix D of LHCSR3 extends into the thylakoid lumen, positioned directly beneath the helix C of CP26 (Fig. 4A). Several acidic residues (E233, E237 and D240) in helix D are potentially protonated and form hydrogen bonds with carbonyl oxygen groups of amino acids located in the lumenal end of helix C and BC loop of CP26 (Figs. 4B). The BC loop of CP26 exhibits a conformation distinct from that of other *C. reinhardtii* Lhcb proteins (Fig. S13A), furthermore, the corresponding region in LHCII participates in trimer formation and therefore inaccessible to LHCSR3 (Fig. S13B). These structural features explain the selective binding of LHCSR3 to CP26 in the C_2_S_2_M_2_L_2_ supercomplexes. The interfacial chlorophyll molecules further stabilize the LHCSR3-CP26 association. Chls 611 and 612 of LHCSR3 are strongly coupled, forming a Chl dimer that is closely connected with the Chl 603-609 pair of CP26, with a shortest Mg-to-Mg distance of 13.1 Å (between Chl 611_LHCSR3_ and Chl 609_CP26_) (Fig. 4A).

**Figure 4.**
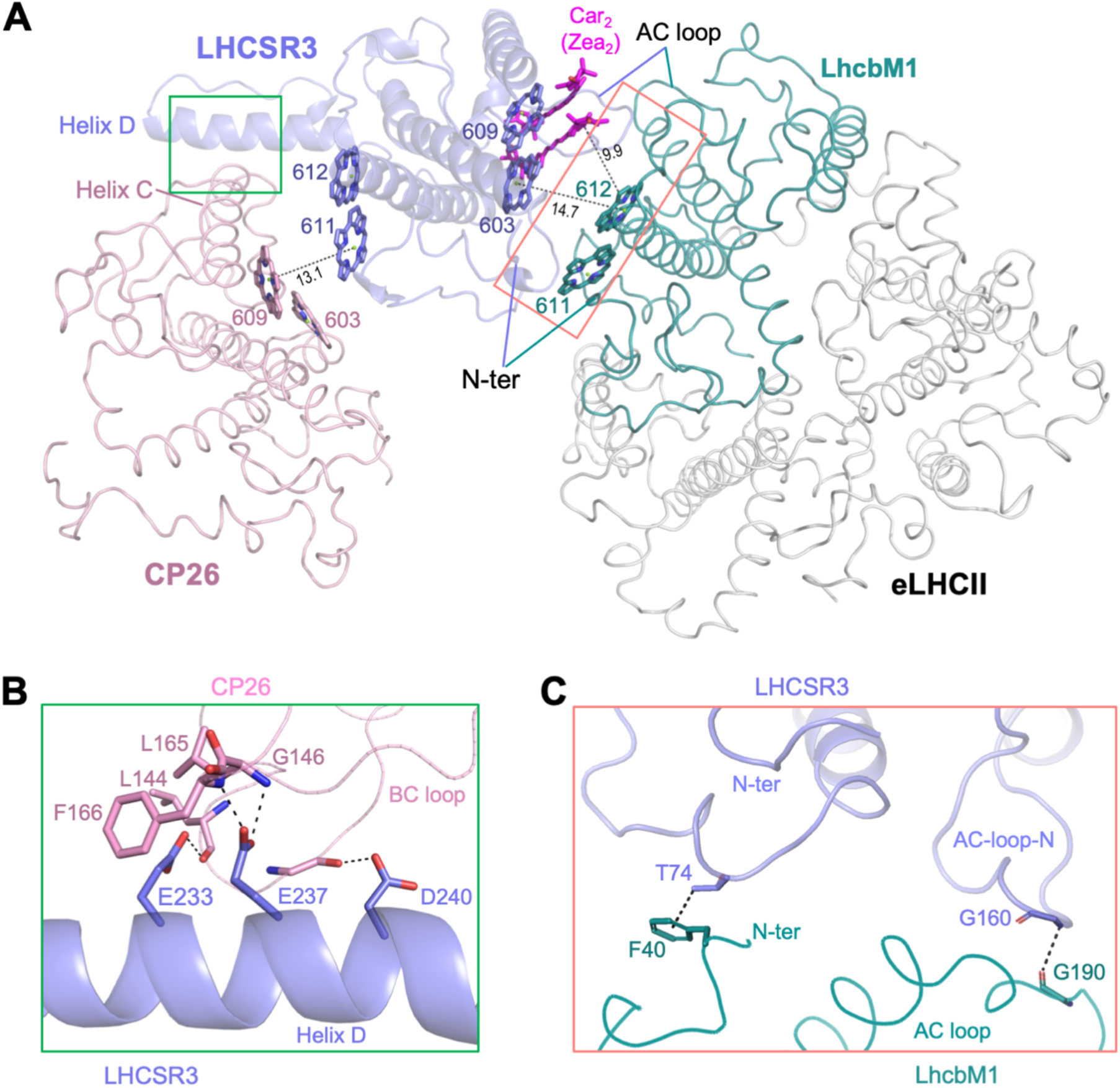
Specific interactions of LHCSR3 with CP26 and eLHCII. **(A)** Stromal side view of the association of LHCSR3 (shown as slate cartoon) with its interacting partners CP26/eLHCII (shown in ribbon mode). The N-terminal and AC loop regions of LHCSR3 and LhcbM1, the helix D of LHCSR3 and helix C of CP26 involved in the interactions are labeled. Pigment molecules involved in the interactions are shown as sticks and labeled. Black dotted lines connect neighboring pigment molecules belonging to different protein subunits, with the distance (Å) labeled adjacent to the line. The phytol tails of chlorophylls are omitted for clarity. **(B)** Enlarged view of the area within the green box in (A). Three acidic residues located in helix D of LHCSR3 form hydrogen bonds (shown as black dashed lines) with the main-chain amino and carbonyl groups of residues from the lumenal end of helix C and BC loop of CP26. **(C)** Enlarged view of the area within the red box in (A), showing detailed interactions between LHCSR3 and LhcbM1 through their N-terminal region (N-ter) and AC loops. Black dashed lines indicate the hydrogen bond and CH–π interactions.

Outside the C_2_S_2_M_2_L_2_ supercomplex, an eLHCII trimer binds to LHCSR3 without forming any direct contact with the C_2_S_2_M_2_L_2_ moiety (Fig. 1B). Based on our cryo-EM map (Fig. S14) and previous biochemical analysis of *C. reinhardtii* LHCII ^49,50^, we tentatively assigned the eLHCII as a LhcbM1-LhcbM2-LhcbM3 heterotrimer, with LhcbM1 oriented toward LHCSR3 (Fig. 4A). The structural observation explains the important role of LhcbM1 in the qE induction ^21^. The interactions between eLHCII and LHCSR3 occur on the stromal side, mediated by the N-terminal and AC loop regions of LhcbM1 and LHCSR3 (Fig. 4A, C). In particular, the AC-loop-N of LHCSR3, with its unique extended conformation (Fig. 3B), plays an important role in binding eLHCII by forming a main-chain hydrogen bond with LhcbM1. Moreover, T74 of LHCSR3 forms a CH–π interaction ^51^ with F40 of LhcbM1 (Fig. 4C), which might be a key residue for LHCSR3 to recognize LhcbM1, as the corresponding residue in LhcbM2 and LhcbM3 is glutamate (Fig. S7A). Furthermore, pigment molecules also contribute to the stability of LHCSR3-eLHCII subcomplex. The Chl 603-609 pair and the carotenoid dimer of LHCSR3 are in close vicinity of the Chl 611-612 pair of LhcbM1 (Fig. 4A), with a shortest Mg-to-Mg distance of 14.7 Å, between Chl 603_LHCSR3_ and Chl 612_LhcbM1_, and the carotenoid dimer approximately 10 Å away from Chl 612_LhcbM1_.

Previous studies have shown that a certain number of LHCII trimers are not tightly bound to the photosystems, but instead form LHCII pools within thylakoid membranes ^49,52,53^. The eLHCII observed in our HL-structure presumably belongs to the LHCII pool and is stabilized by binding to LHCSR3 under high-light conditions. Consistent with this suggestion, we observed extra densities surrounding the C_2_S_2_M_2_L_2_ complex in both HL- and LL-reconstructions when the maps were low-pass filtered to 15 Å (Fig. S15). These densities presumably correspond to extra LHCII trimers in the pool, loosely and heterogeneously associated with the PSII-LHCII supercomplexes, and some of which are positioned approximately 20-40 Å away from the C_2_S_2_M_2_L_2_ supercomplex. Therefore, these extra LHCIIs may transfer excitation energy to PSII under low-light, albeit less efficiently than those closely bound LHCIIs (S-, M- and L-LHCIIs). Under high-light conditions, the regulation of light harvesting and energy transfer may also target these extra LHCIIs through the specific binding of LHCSR3 to eLHCII.

### Two unique pigment clusters in LHCSR3

Under high-light conditions, LHCSR3 may act as a central qE quencher, directly dissipating excess excitation energy from the PSII-LHCII complex and eLHCII via its carried pigment molecules. Quencher pigments typically include energetically coupled chlorophyll-carotenoid (Chl-Car) clusters and Chl dimers ^54,55^. While all eight chlorophylls and two stably-bound carotenoids in LHCSR3 are conserved in other LHC proteins (Fig. S10B, C), their orientations differ slightly from their counterparts in LhcbM1 and Lhcx6_1 (Fig. 5A). The Chl 611 in LHCSR3 exhibits the most pronounced difference, with its porphyrin ring pivoted by approximately 20 degrees relative to that in LhcbM1 (Fig. 5B), presumably because LHCSR3 lacks the PG molecule that coordinates Chl 611 in other Lhcb proteins (Fig. S11). Similarly, the porphyrin ring of Chl 603 in LHCSR3 also pivots by ∼10 degrees relative to the corresponding one in LhcbM1 (Fig. 5C). As a result, the Chl 611-612 and 603-609 pairs in LHCSR3 have nearly parallel and partially overlapping porphyrin rings with short ring-ring distances (< 4 Å), thereby forming two strongly coupled Chl dimers, similar to the special pair in the reaction center of PSII/PSI. The parallel arrangement of the two Chl dimers was also observed in Lhcx6_1, albeit they are slightly less parallel than those in LHCSR3 (Fig. 5B, C). Notably, these two Chl dimers are distributed on opposite sides of LHCSR3 (Fig. 5A, D), facing CP26 and eLHCII respectively, and directly mediating interactions of LHCSR3 with these two antenna components (Fig. 4A). Such an arrangement and location may enable these two Chl dimers to act as energy traps to capture the excitation energy absorbed by other antenna proteins and/or as quenchers to dissipate excess energy ^54,56^.

**Figure 5.**
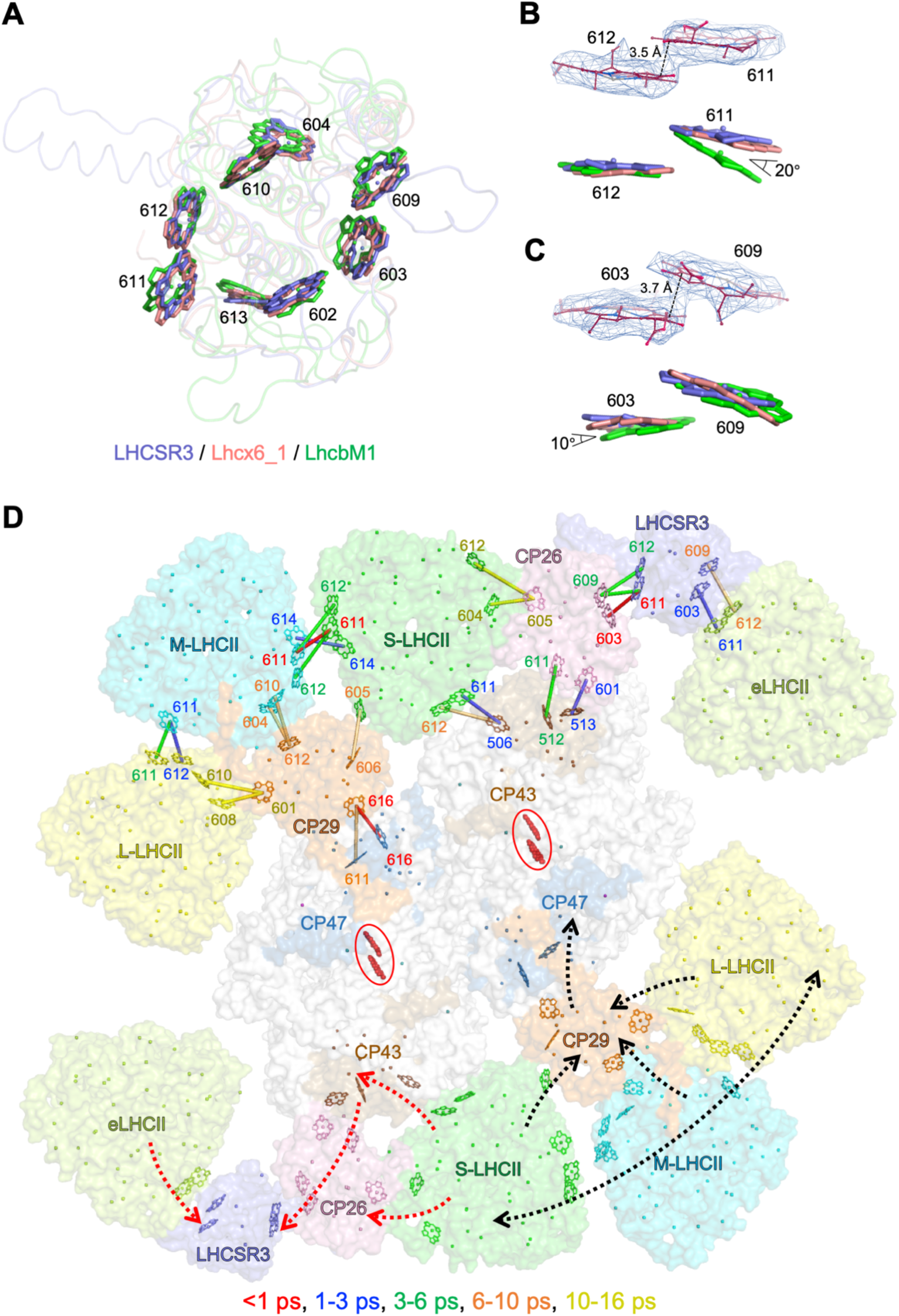
Chlorophyll arrangement in LHCSR3 and the potential energy transfer in the HL_PSII-LHCSR supercomplex. **(A)** Comparison of chlorophyll arrangement and orientation in LHCSR3 (slate), Lhcx6_1 (salmon) and LhcbM1 (green), viewed from the stromal side. Proteins are shown as transparent ribbon. Eight conserved chlorophyll molecules are shown as sticks and labeled. Other chlorophyll molecules in Lhcx6_1 and LhcbM1 are omitted for clarity. **(B, C)** Cryo-EM densities of Chl dimers 611-612 (B) and 603-609 (C) in LHCSR3, and structural comparison of the two Chl dimers in LHCSR3 (slate), Lhcx6_1 (salmon) and LhcbM1 (green). **(D)** Rapid energy transfer pathways in the HL_PSII-LHCSR supercomplex based on the FRET calculations, viewed from the stromal side. Proteins are shown in surface mode, with PSII core subunits CP43, CP47 and LHC proteins shown in different colors and labeled. Other subunits are shown in white. Chlorophyll molecules are shown as spheres at their central-Mg positions, with chlorophylls involved in the rapid EETs between subunits also shown as sticks and labeled in the upper half of the supercomplex. The two chlorophyll molecules in D1 and D2 subunits forming the special pair (P680) are shown in stick-ball mode in red, and highlighted by red circles. In the upper half of the supercomplex, lines connecting two neighboring chlorophylls represent the rapid EET pathways, and the colors of lines indicate different lifetimes, as annotated in the bottom. The chlorophyll labels are colored the same as the lines connecting the chlorophyll molecules. For clarity, only the fastest EET (two to four shortest lifetimes between different protein subunits) are shown, and all chlorophyll pairs with rapid EET (lifetime < 20 ps) are listed in Table S3. In the lower half of the supercomplex, black and red dashed lines represent the proposed energy transfer and energy quenching process, respectively.

We further calculated the Förster Resonance Energy Transfer (FRET) rate between chlorophyll molecules belonging to adjacent subunits in the HL_PSII-LHCSR complex based on our structure (Fig. 5D, Table S3). Consistent with our structural analysis, the calculations indicate that Chl dimer 611-612/603-609 in LHCSR3 play a crucial role in mediating rapid excitation energy transfer (EET) between LHCSR3 and CP26/eLHCII. In particular, EET between LHCSR3 and CP26 is extremely fast, occurring on a sub-picosecond (ps) timescale between Chl 611_LHCSR3_ and Chl 603_CP26_, while the rapid energy equilibration between LHCSR3 and eLHCII is achieved in less than 3 ps via Chl 603_LHCSR3_ and Chl 611_eLHCII_. Furthermore, CP26 and S-LHCII also exhibit fast EET with CP43 in the picosecond range, with the shortest lifetimes of approximately 2-3 ps. These data suggest that LHCSR3 can efficiently trap the excitation energy from CP26 (and even CP43 and S-LHCII) and eLHCII (and LHCII pool) via its two Chl dimers.

LHCSR3 has been suggested to be active in the Chl-Car charge-transfer quenching mechanism ^57^, and a signal similar to that of Lut radical cation was detected in the reconstituted LHCSR3 *in vitro* ^12,44^. Moreover, a recent study revealed that an ultrafast energy transfer from Chl *a* to an adjacent Lut molecule plays a role in the qE function of LHCSR3 ^58^. In LHCSR3 structure, the two carotenoid molecules at L1 and L2 sites are tightly bound to Chl 611-612 and 603-609 dimers, respectively. In addition, Chl 603-609 is further associated with the carotenoid (Zea) dimer, resulting in the formation of two Chl dimer-Car clusters, namely 611-612-Lut and 603-609-Vio-Car_2_ (603-609-Vio-Zea_2_) in LHCSR3, with distances of approximately 4 Å between adjacent pigments within each cluster (Fig. 6A). Lutein and zeaxanthin are two types of carotenoids capable of quenching singlet and triplet chlorophylls ^12,54,59^, and Zea dimers have been previously suggested to be able to quench Chl *a* fluorescence ^36^, thus these two Chl dimer-Car clusters in LHCSR3 likely play a critical role in qE. In addition to the Chl dimer, Lut and Vio (the L1 and L2 carotenoids) are also closely associated with Chl 610/613 and Chl 602/604 at the stromal/lumenal side, respectively (Fig. 6B, C). Furthermore, we observed one putative carotenoid molecule in LHCSR3 located adjacent to Chls 611 and 613 (Fig. S12F), which corresponds to a fucoxanthin (Fx301) in Lhcx6_1, showing the same binding position and the same interaction pattern ^43^. As a result, each chlorophyll molecule in LHCSR3 is closely coupled to at least one carotenoid molecule, and all its pigment molecules can be divided into two clusters, namely the L1-cluster comprising Chl 611-612 dimer, Lut, Chls 610 and 613 (and the putative carotenoid), and L2-cluster comprising Chl 603-609 dimer, Vio, the carotenoid (Zea) dimer, Chls 602 and 604 (Fig. 6B, C). Interestingly, we found that both LHCSR3 and Lhcx6_1 possess phenylalanine residues in helix D and BC-loop-N regions, which are presumably located close to Chls 613 and 604 under low light conditions (Fig. 6B, C). After high light illumination, the protonation-induced conformational changes in the lumenal regions of LHCSR3 may affect Chls 604 and 613 through these phenylalanine residues, and further transmit to carotenoid molecules as well as the stromal side chlorophylls.

**Figure 6.**
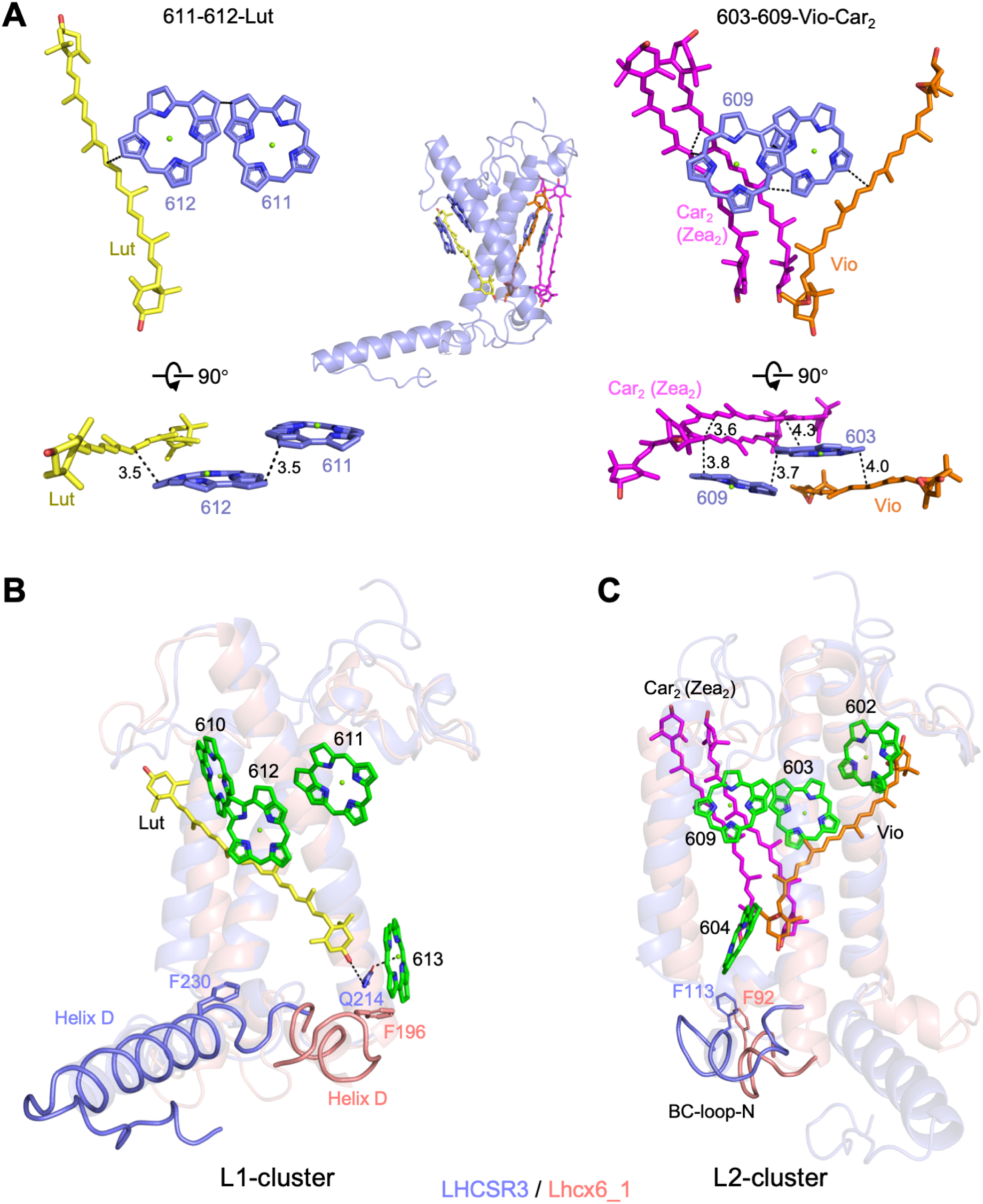
Two distinct pigment clusters in LHCSR3. **(A)** Distribution and arrangement of 611-612-Lut (left) and 603-609-Vio-Car_2_ (right) clusters. Color codes are the same as in Fig. 2A. The shortest distances between neighboring pigment molecules are indicated by black dashed lines, with distances (Å) labeled adjacent to the lines. **(B)** The L1-cluster in LHCSR3, with Lhcx6_1 apoprotein (salmon) superimposed on LHCSR3 (slate). Proteins are shown as transparent cartoon, with the C-terminal regions including helix D also shown as ribbons. Pigments in L1-cluster of LHCSR3 are shown as sticks and labeled. The conserved residue Q214 in LHCSR3, which coordinates Chl 613 and forms a hydrogen bond with Lut, is shown as sticks and labeled. The Chl 613 is located at the lumenal side and might be close to helix D of LHCSR3 under low-light conditions, as represented by helix D of Lhcx6_1. Both LHCSR3 and Lhcx6_1 harbors a phenylalanine (shown as lines) in helix D, with F196 in Lhcx6_1 located close to Chl 613, whereas F230 in LHCSR3 being distant from Chl 613 due to the different orientation of the helix D. **(C)** The L2-cluster in LHCSR3, with Lhcx6_1 apoprotein (salmon) superimposed on LHCSR3 (slate). Proteins are shown as transparent cartoon, with the BC-loop-N regions also shown as ribbons. Pigments in L2-cluster of LHCSR3 are shown as sticks and labeled. Chl 604 is located at the lumenal side and close to BC-loop-N of LHCSR3. Both LHCSR3 and Lhcx6_1 harbors a phenylalanine in BC-loop-N (F113 in LHCSR3 and F92 in Lhcx6_1), which is located close to Chl 604 and shown as lines.

## DISCUSSION

Despite extensive researches on LHCSR over a decade, the molecular mechanisms of qE involving LHCSR are still unclear, and structures of LHCSR and its complex with the PSII remain unknown. Our previous attempts to purify PSII-LHCSR complex and solve its structure using single particle cryo-EM proved unsuccessful, likely due to the weak binding affinity of LHCSR3 to PSII and its low stoichiometry. Here, enabled by our newly developed cryo-LEP-FIB technique in combination with the isSPA approach, we successfully solved the *in situ* structures of *C. reinhardtii* HL_PSII-LHCSR and LL_PSII complexes at near-atomic resolutions (Fig. 1). We found that under high-light conditions, LHCSR3 exists as a monomer and binds to the C_2_S_2_M_2_L_2_ supercomplex at a specific position adjacent to CP26, and also interacts with an eLHCII presumably originating from the LHCII pool. In comparison, LHCSR3 does not bind to the PSII-LHCII under low-light conditions (Fig. S4). These structures likely represent the native conformations of PSII-LHCII in quenched and unquenched states, respectively. In addition, our results suggest that *in vivo*, the C_2_S_2_M_2_L_2_ type is the most abundant PSII-LHCII form in *C. reinhardtii* even under high-light conditions, consistent with a recent study indicating that qE does not lead to the structural disassembly of PSII complex^60^.

Comparison of LHCSR3 with diatom Lhcx6_1 (Fig. 3A-C), the only LHCSR homolog with a known structure ^40–43^, suggested potential conformational rearrangements in LHCSR3 in response to thylakoid lumen acidification. Presumably, LHCSR3 adopts a fold similar to Lhcx6_1 under low light conditions, exhibiting the same orientation of helix D. The HL-induced decrease in lumen pH leads to the protonation of several acidic residues located in the lumenal regions, including the three qE-crucial residues (D117, E221 and E224). The protonation event may induce unwinding of the N-terminus of helix D and its subsequent large-scale rotation, which is critical for mediating the LHCSR3-CP26 association (Fig. 4B). Thus the lack of LHCSR3-bound PSII in LL-cells may be because of the unfavorable conformation of LHCSR3 in addition to its low abundance. Furthermore, LHCSR3 may recruit and interact with a Zea dimer through its BC loop and AC-loop-N regions (Fig. S12D), which may stabilize the AC-loop-N conformation, enabling it to bind to eLHCII (Fig. 4A, C). In addition, the Zea dimer and other chlorophyll molecules may further reinforce the stability of the LHCSR3-eLHCII subcomplex. These structural observations suggest that the simultaneous binding of eLHCII and the Zea dimer to the AC-loop-N of LHCSR3 may mutually stabilize each other.

The recruitment of LHCSR3 to the position in association with CP26 and eLHCII is likely essential for establishing efficient EET pathways between LHCSR3 and CP26/eLHCII. Consistent with this suggestion, our FRET rate calculations indicate that EET between LHCSR3 and CP26/eLHCII is highly effective, with Chl 611-612 and 603-609 dimers in LHCSR3 serving as the primary mediators (Fig. 5D, Table S3). The activated LHCSR3 dissipates excess energy transferred from PSII-LHCII and eLHCII, a process presumably achieved through the pigment rearrangement caused by the conformational changes in lumenal regions of LHCSR3. Previous studies have shown that removing Chl 613 by mutating its coordinating residue reduces the quenching efficiency of LHCSR3, suggesting that Chl 613 is likely involved in thermal dissipation ^35,61^. The Chl 613 is located close to the Lut molecule, and its coordinating residue Q214 forms a hydrogen bond with the hydroxyl group of the lumenal ring of Lut (Fig. 6B). However, the weak density of Chl 613 (Fig. 2E) indicates its loose binding to LHCSR3 under high-light, which is potentially due to the rotation of helix D. Under low-light conditions, Chl 613 is presumably stabilized by helix D in an orientation similar to that of Lhcx6_1 (Fig. 6B). Based on these results, we propose that the pivot of helix D induced by lumen acidification may destabilize Chl 613, thereby affecting Lut and leading to conformational rearrangements within the L1-cluster and subsequent activation of quenching. Removal of Chl 613 may have an impact on helix D and the L1-cluster, thus affecting the quenching efficiency of LHCSR3. Similarly, Vio in the L2-cluster is closely associated with Chl 604 via its lumenal ring, and Chl 604 is located adjacent to the BC-loop-N (Fig. 6C), thus the L2-cluster may be affected by lumenal conformational changes transmitted through Chl 604. Therefore, the two clusters may respond to the lumen acidification and function in the pH-dependent quenching ^12,57,58^. Furthermore, the L2-cluster contains a putative Zea dimer (Fig. 6C), and other Zea molecules might also assemble with LHCSR3 at its periphery, but exhibit high flexibility and low occupancy (Fig. S12F). Residues that interact with these putative Zea molecules are highly conserved across LHCSR proteins from different species (Figs. S11B, S12D), underscoring their critical role in LHCSR function. These putative Zea molecules may account for the previously reported Zea-dependent quenching of LHCSR3 ^20^. Furthermore, the interfacial locations of these Zea molecules suggest that they play an important role in stabilizing the HL_PSII-LHCSR complex, and may also contribute to NPQ by improving the energetic connectivity between quenchers and antenna complexes, as suggested in a recent report ^55^.

Based on our structural analysis and previously reported data ^11,12,15,25,48^, we hereby propose a mechanistic model for the quenching process in the HL_PSII-LHCSR complex (Fig. 7). Under high-light conditions, LHCSR3 expression is upregulated, leading to the increased occupancy of LHCSR3 nearby the PSII-LHCII supercomplexes. Moreover, the decrease of lumen pH results in the protonation of lumenal acidic residues in LHCSR3, including D117, E221 and E224. Protonation of these residues induces conformational changes in these lumenal regions, particularly helix D and BC-loop-N, and the subsequent rearrangements of pigment clusters within LHCSR3. These changes may drive the transition of LHCSR3 from its unquenched to quenched state, enabling it to interact directly with C_2_S_2_M_2_L_2_ supercomplex via CP26 to dissipate energy transferred to PSII, and to stabilize the eLHCII from the LHCII pool, thereby expanding the quenched region of light-harvesting antennae ^55,62^. Furthermore, high-light also results in the conversion of Vio to Zea in the lipid phase ^63,64^. LHCSR3 may recruit these newly synthesized Zea molecules at the LHCSR3-eLHCII and LHCSR3-CP26 interfaces, where they stabilize the HL_PSII-LHCSR complex and participate in thermal dissipation. The two pigment clusters distributed on opposite sides of LHCSR3 may serve as two quenching sites targeting different antenna components. The Chl 611-612 dimer and associated carotenoid molecule(s) in the L1-cluster face CP26 and may thus dissipate energy transferred from CP26 (and potentially other parts of PSII-LHCII through CP26, such as CP43 and S-LHCII). The Chl 603-609 dimer and carotenoids in the L2-cluster are close to eLHCII and could attenuate energy transmitted from the LHCII pool to PSII by quenching the energy transferred from eLHCII. Therefore, the formation of the HL_PSII-LHCSR complex enables LHCSR3 to quench the excitation energy absorbed by PSII-LHCII itself and also that transferred from the LHCII pool, thereby protecting PSII from photodamage. Recent studies have demonstrated that accelerating the NPQ process can increase photosynthetic productivity in tobacco and soybean ^65,66^. Therefore, understanding the mechanisms of qE is essential for precisely regulating this process to increase biomass. Our research represents a key step forward, providing an important foundation for further improving microalgae productivity under fluctuating light conditions.

**Figure 7.**
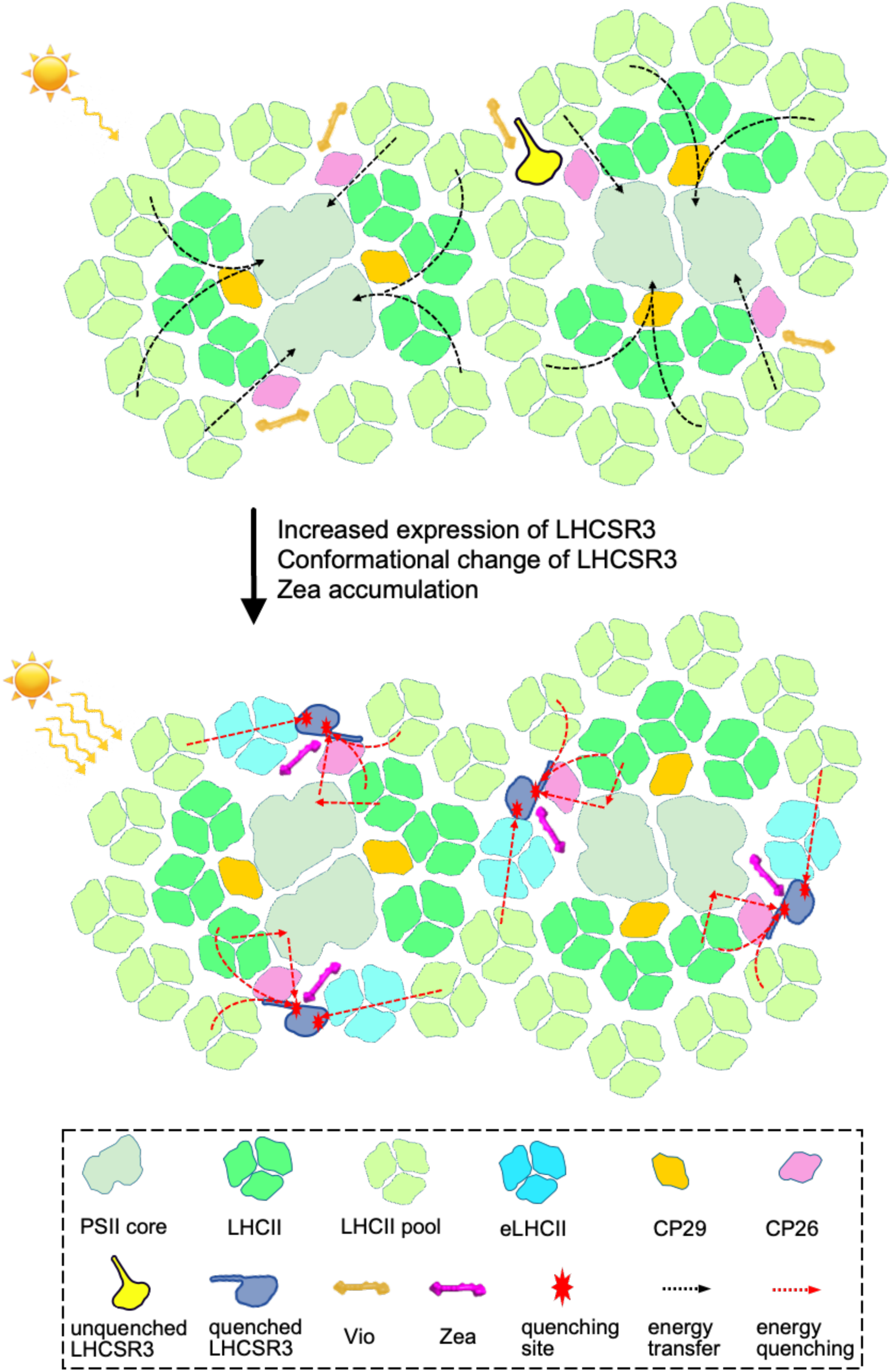
Proposed model of LHCSR3 quenching excess energy from PSII-LHCII and LHCII pool.

## METHODS

### *C. reinhardtii* culture conditions

To obtain LL-cells, the *C. reinhardtii* CC124 cells were cultured in TAP medium at 23 °C with continuous shaking at 130 rpm under constant low light-intensity (30-40 μmol photon m^-2^ s^-1^). To obtain HL-cells, the LL-cells were first treated under high-light illumination (500 μmol photons m^-2^ s^-1^) for 12 hours. Both LL-and HL-cells were harvested by centrifugation at 1,000 g for 5 min, and then resuspended in fresh TAP medium to a concentration of 0.01 mg ml^-1^ (in Chl) for cryo-EM specimen preparation. Since the 5-min centrifugation may cause qE relaxation in HL-cells, the centrifuged HL-cells were further exposed to HL conditions for 2 hours before sample preparation to ensure that the HL-cells are in the quenched state.

### Sample preparation and Cryo-FIB milling

Approximately 3 µL aliquot of cells (0.01 mg ml^-1^ in Chl) was applied onto a glow-discharged copper Quantifoil R 1.2/1.3 200 mesh grid. The back-side blotting was performed for 4-6 s at 16°C and 70% chamber humidity, and the grid was subsequently plunge-frozen in a 1:1 mixture of liquid ethane and propane using a Leica EM GP (Leica Microsystems). The HL illumination was maintained throughout the grid preparation process.

Cryo-lamellae were prepared from the frozen grids using cryo-focused ion beam (cryo-FIB) milling in an Aquilos 2 FIB-SEM microscope (Thermo Fisher Scientific). Prior to milling, the sample surface was sequentially sputter-coated with metallic platinum for 15 s, organo-platinum for 60 s via the gas injection system (GIS), and then re-coated with metallic platinum for an additional 15s. Rough milling was performed using Thermo Fisher’s AutoTEM Cryo 2.4 software, progressively reducing the ion beam current from 1.0 nA to 50 pA at 30 keV and a milling angle of 10°, achieving an approximate thickness of 300 nm. Finally, to minimize damage, low-energy polishing was conducted at 8 keV and 75 pA, further thinning the lamella to less than 150 nm.

### Data collection and image processing by isSPA

Approximately 50 cryo-lamellae from HL-cells and 12 cryo-lamellae from LL-cells were collected on an FEI Titan Krios G3 microscope (Thermo Fisher Scientific) equipped with a Gatan K2 Summit direct electron detector camera (Gatan Inc.). Prior to collection, an initial stage tilt offset of 10° was applied to compensate for the FIB milling angle, ensuring the lamellae remained perpendicular to the electron beam during imaging, as previously described ^29,67^. Data collection for both datasets was conducted using SerialEM with the beam-image shift method ^68^. Images were recorded at a nominal magnification of 105,000×, corresponding to a calibrated physical pixel size of 1.32 Å/pixel. The total exposure dose was 40 e^-^Å^-^², fractionated over 40 frames with a total exposure time of 6 s. The nominal defocus range was set between -0.8 and -1.2 μm.

Motion correction and dose weighting were done by MotionCor2 ^69^ with a binning factor of 2. The CTF parameters of micrographs were estimated by CTFFIND4 ^70^ in RELION3 ^71^. Then 4,891 the micrographs of thylakoid membranes in HL condition were selected for particle matching by isSPA ^28,29^, using the dimeric PSII core (C_2_) region of *in situ* PSII map reconstructed based on EMD-9956 ^32^ as the template (details see companion paper). The template was projected at 3° angular intervals, generating 4,582 2D projections (excluding in-plane rotations). Particle matching was performed with overlapping parameter *n* as 3 and a frequency range of 1/400 Å^-1^ to 1/6 Å^-1^ yielding 1,969,163 particles with a score threshold of 6.5. Particles were extracted with 2× binning in a box size of 400 pixels, followed by alignment-free 3D classification in RELION3. Approximately 35.8% of particles, containing S-, M-, and L-type LHCIIs, were selected for another round of alignment-free 3D classification. A total of 260,035 particles were then used for local refinement with unbinned (full-size) extraction, resulting in a 4.0 Å reconstruction of the C_2_S_2_M_2_L_2_-type PSII supercomplex. To further reduce potential bias, a sorting algorithm ^28^ in isSPA was applied, and particles below a score threshold of 0.0276 were discarded, removing 57.2% of the dataset. Another round of 3D classification was then performed in cryoSPARC ^72^ using a mask covering the LHCSR3 and the extra LHCII (LHCSR-eLHCII) after C2 symmetry expansion. From this, 57.9% of particles containing LHCSR-extra densities were selected for OpticsGroup CTF refinement in RELION3, yielding an overall resolution of 3.1 Å. Local refinements of CP26 and S-LHCII regions (CP26-S) yielded a 3.0 Å map, while refinements of CP29, M- and L-LHCII (CP29-M-L) resulted in a 3.0 Å map. To further improve the resolution of the LHCSR-eLHCII region, another round of 3D classification was conducted in cryoSPARC using the same LHCSR-eLHCII mask. Two similar classes, comprising 112,861 particles with more continuous densities in this region, were selected for local refinement with a gradually shrinking mask, ultimately achieving a local resolution of 3.8 Å (Fig. S2).

892 micrographs collected under low-light (LL) conditions were processed using a similar workflow (detailed data processing is presented in the companion paper), resulting in a 3.4 Å overall resolution of the C_2_S_2_M_2_L_2_-type PSII structure, based on a final particle set of 114,764 (after C2 symmetry expansion). The local resolution reached 3.3 Å in the CP26 and S-LHCII regions (CP26-S), and 3.4 Å in the CP29, M- and L-LHCII regions (CP29-M-L) (Fig. S3).

### Model building and refinement

For model building of HL_PSII-LHCSR, the cryo-EM structure of *C. reinhardtii* C_2_S_2_M_2_L_2_-type PSII (PDB code: 6KAD)^32^ was manually fitted into the 3.1 Å *in situ* cryo-EM map using UCSF Chimera ^73^. Several protein subunits missing from the C_2_S_2_M_2_L_2_ structure (PsbP, PsbQ, and Psb30) were obtained from the C_2_S_2_-type PSII structure (PDB code: 6KAC) and manually fitted into the map. PsbR was automatically built using CryoAtom ^74^ and the sequence was subsequently mutated to match *C. reinhardtii* PsbR sequence. For LHCSR3 and PsbY, the AlphaFold-predicted models were fitted into the map, and the C-terminal region of LHCSR3 was manually adjusted to align the cryo-EM density. The *de novo* model building was performed based on the cryo-EM map for the previously unidentified protein Psb12 and PsbTn, since the PsbTn sequence was not identified in the UniProt database and our searches using sequences of its plant homologs were unsuccessful. Subsequently, the partial sequences derived from the manually built models of these two proteins were used to search for their full-length sequences in the UniProt database, and identified two uncharacterized proteins, XP_001690252.1 and XP_042921420.1, as *C. reinhardtii* Psb12 and PsbTn, respectively. The identification of each individual LhcbM protein was based on the best match of the specific amino acids in sequences and the cryo-EM densities.

Real-space refinement utilizing the Phenix program (v. 1.21.2) ^75^ and manual adjustment using COOT ^76^ were then performed interactively. The geometries of the structural model were evaluated using MolProbity ^77^, and the detailed information of data collection and structure refinement are summarized in Table S1.

### Identification and quantification of pigments by HPLC

LL- and HL-cells were harvested by centrifugation and resuspended in TAP medium to a final concentration of 1.0 mg ml^-1^ (in Chl). The PSI-LHCI and PSII-LHCII supercomplexes were purified from HL-cells through sucrose density gradient ultracentrifugation according to previous reports ^32,78^. For pigments extraction, a mixture of 20 μl cells (or purified samples of PSI-LHCI or PSII-LHCII complexes) and 80 μl acetone was vortexed thoroughly and centrifuged twice at 17,500 g for 5 minutes at 4°C. The supernatant was loaded on a C-18 reversed-phase column (Alltech Allsphereg ODS-2), and pigments were eluted following the previously established protocol ^79^. Individual pigments were identified by the absorption spectrum and retention time of each peak. The HPLC system comprised a Hitachi L2130 separation module equipped with a Hitachi L2450 diode array detector.

### NPQ and Fv/Fm measurement

*C. reinhardtii* cells acclimated to HL or LL conditions were harvested by centrifugation at 1,000 g for 5 min, and resuspended in fresh TAP medium to a final concentration of ∼0.03 mg ml^-1^ in Chl. Cells were dark-adapted for 30 minutes at room temperature prior to NPQ measurement. Chlorophyll fluorescence was recorded with a Dual-PAM-100 (Walz) instrument. The measuring light intensity was set to <1 μmol photons m^−2^ s^−1^, the actinic light was set to 1,400 μmol photons m^−2^ s^−1^, and the saturating pulses were set to 6,000 μmol photons m^−2^ s^−1^. Fo and Fm were determined after dark adaptation, and Fm′ was determined during illumination with actinic light. NPQ was calculated as (Fm – Fm′)/Fm′ according to an earlier report ^3^. All measurements were performed at room temperature with gentle stirring, and repeated three times using independent biological replicates. Fv/Fm was used to evaluate the maximum photochemical efficiency of PSII and was calculated as (Fm–Fo)/Fm.

### Immunoblot analysis of LHCSR

Equal amount (1 μg Chls) of HL-cells and LL-cells were boiled for 5 min after adding loading buffer, then separated by sodium dodecylsulfate polyacrylamide gel electrophoresis (SDS-PAGE) on a 12% polyacrylamide gel. LHCSR3 antibody was purchased from Agrisera (1:1,000 dilutions), and the secondary antibody ligated to horse radish peroxidase (1:2,000 dilution) was applied. Western blot analysis was performed using EasySee Western Blot Kit from TransGen Biotech on the PVDF membrane, and images were recorded on CLiNX Imaging System.

### FRET calculation

Förster resonance energy transfer (FRET) rate constants (*k*_FRET_) were calculated according to the classical FRET theory ^80^ using the expression *k*_FRET_=(*CK*^2^)/(*n*^4^*R*^6^), where R denotes the distance between the central Mg atoms of the donor and acceptor chlorophylls, n is the refractive index of the medium (set to 1.55), C is a constant derived from the spectral overlap integral between the two chlorophylls, as estimated in reference ^81^, the C values is 32.5 (Chl a to Chl a), 1.11 (Chl a to Chl b), 9.61 (Chl b to Chl a) and 14.45 (Chl b to Chl b), respectively. K^2^ represents the dipole orientation factor, is defined as 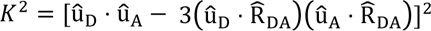, with 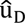 and 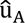 being the unit vectors of the transition dipoles for the donor and acceptor chlorophylls, respectively. These dipole vectors were derived from the coordinates of NB and ND atoms, 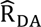 is the unit vector along the direction connecting the Mg atom of the donor to the acceptor chlorophylls. All FRET rates were computed using Kim’s algorithm on the Python platform (Python v.3.8) and are available at https://doi.org/10.5281/zenodo.3250649.

## ACKNOWLEDGMENTS

Cryo-EM datasets were collected at the Center for Biological Imaging (CBI), Core Facilities for Protein Science at the Institute of Biophysics (IBP), Chinese Academy of Sciences (CAS). We thank staff members at the CBI (IBP, CAS) for technical support in data collection. We are grateful to Profs. Minrui Fan, Xiaowei Pan and Jean-David Rochaix for in-depth discussions.

## AUTHOR CONTRIBUTIONS

M.L. and X.Z. conceived and supervised the project; X.S. performed the sample preparation and characterization, model building and refinement; C.W. performed data collection and processing; S.C. helped in sample preparation and computational modeling; M.L., X.S. analyzed the structures; M.L., X.S., C.W. and Z.L. wrote the manuscript; all authors discussed and commented on the results and the manuscript.

